# Extend Mixed Models to Multi-layer Neural Networks for Genomic Prediction Including Intermediate Omics Data

**DOI:** 10.1101/2021.12.10.472187

**Authors:** Tianjing Zhao, Jian Zeng, Hao Cheng

## Abstract

With the growing amount and diversity of intermediate omics data complementary to genomics (e.g., DNA methylation, gene expression, and protein abundance), there is a need to develop methods to incorporate intermediate omics data into conventional genomic evaluation. The omics data helps decode the multiple layers of regulation from genotypes to phenotypes, thus forms a connected multi-layer network naturally. We developed a new method named NN-LMM to model the multiple layers of regulation from genotypes to intermediate omics features, then to phenotypes, by extending conventional linear mixed models (“LMM”) to multi-layer artificial neural networks (“NN”). NN-LMM incorporates intermediate omics features by adding middle layers between genotypes and phenotypes. Linear mixed models (e.g., pedigree-based BLUP, GBLUP, Bayesian Alphabet, single-step GBLUP, or single-step Bayesian Alphabet) can be used to sample marker effects or genetic values on intermediate omics features, and activation functions in neural networks are used to capture the nonlinear relationships between intermediate omics features and phenotypes. NN-LMM had significantly better prediction performance than the recently proposed single-step approach for genomic prediction with intermediate omics data. Compared to the single-step approach, NN-LMM can handle various patterns of missing omics measures, and allows nonlinear relationships between intermediate omics features and phenotypes. NN-LMM has been implemented in an open-source package called “JWAS”.

## Introduction

The advances in high-throughput sequencing technology provide growing amount and diversity of multi-omics data complementary to genomics (e.g., DNA methylation, gene expression, and protein abundance). As demonstrated in Figure 1, the effects of genotypes on phenotypes can be mediated by multiple layers of omics features through mechanisms such as regulatory cascades from epigenome, to transcriptome, and to proteome (Ritchie *et al*. 2015; Sun and Hu 2016; Wu *et al*. 2018). This multi-layer regulation works as a unified system to connect genome variations to the trait, and the relationships between different layers can be complex with interactions and nonlinear relationships (Kitano 2002; Green *et al*. 2017; Devijver *et al*. 2017; Green *et al*. 2019). For example, Green *et al*. (2017) observed that the relationship between gene expression level and phenotype was non-linear, which was approximated by a generalised logistic function.

**Figure 1.**
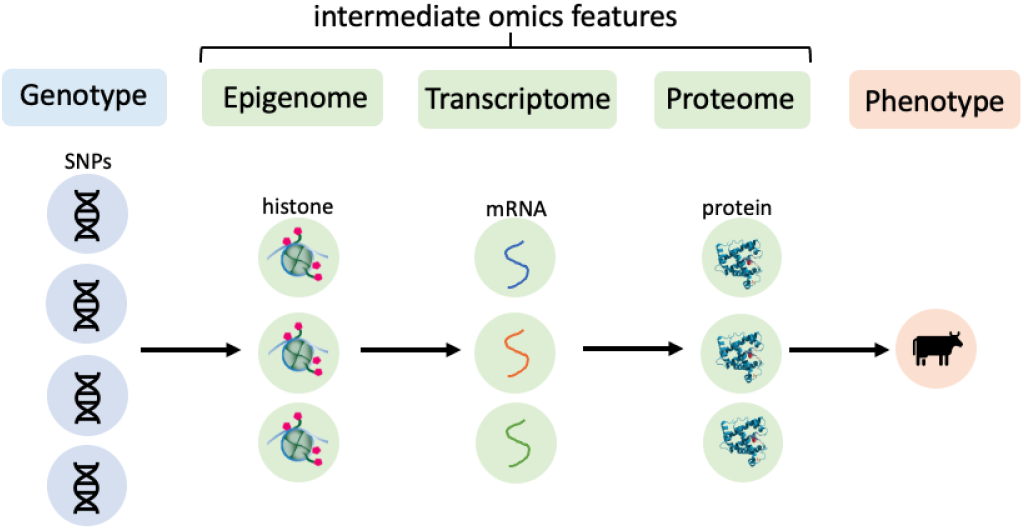
An example of multiple layers of regulation between genotypes to phenotypes. The DNA sequence variations may affect the phenotypes through epigenome, transcriptome, and proteome levels, and the relationships between different layers may be complex with interactions and nonlinear relationships. This unified multi-layer regulation system forms a connected network naturally.

In genotypes-to-phenotypes studies such as genomic prediction (Meuwissen *et al*. 2001; Hayes *et al*. 2009a; Heffner *et al*. 2009; Hickey *et al*. 2017) and genome-wide association studies (GWAS) (Ozaki *et al*. 2002; Visscher *et al*. 2012, 2017; Atwell *et al*. 2010; Korte and Farlow 2013), incorporating intermediate multi-omics data has facilitated our understanding of the relationship between genotypes and phenotypes (Qian *et al*. 2019; Ritchie *et al*. 2015; Christensen *et al*. 2021; Riedelsheimer *et al*. 2012). To incorporate intermediate omics data (e.g., gene expression levels) between genotypes and phenotypes for association studies, approaches such as multi-staged analysis (Ritchie et al. 2015) and transcriptome-wide association studies (Gamazon et al. 2015; Gusev *et al*. 2016; Wainberg *et al*. 2019) were proposed. In these approaches, two linear models were used to describe the relationship between phenotypes and gene expression levels, and the relationship between gene expression levels and genotypes. Such system of two linear models has also been developed recently for genomic evaluation (Weishaar *et al*. 2020; Christensen *et al*. 2021), and further extended for genomic prediction using incomplete omics data (Christensen *et al*. 2021).

In practice, there often exist missing measures in the intermediate omics data, because the omics features are not measured for all individuals who have phenotypes, or different omics features are measured in different experiments. Thus, there is a need to develop methods to model the unified system of multi-layer regulation from genome to intermediate omics features then to the phenotypic trait, with a capability of dealing with missing omics data. Christensen *et al*. (2021) proposed a method to include intermediate omics features into genetic evaluation, in which two linear mixed model equations are required. The first model describes how intermediate omics features affect phenotypes, and the second model describes how genotypes affect intermediate omics features. When omics data for some individuals are completely missing, single-step approach is used to construct the relationship matrix for individuals with all omics features measured and those having no omics data. This is analogous to the popular genomic single-step approach (Legarra *et al*. 2009; Christensen and Lund 2010; Legarra *et al*. 2014) that combines information from both genotyped and non-genotyped relatives in genetic evaluation. As we will show in this paper, this approach for genomic prediction using intermediate omics data is not able to incorporate omics data that are partially missing for some individuals, assume linear relationships between gene expressions and phenotypes, and may give suboptimal results even when the underlying relationship is linear.

As illustrated in Figure 1, the multi-layer regulatory system forms a connected network naturally, thus the architecture of artificial neural networks can be considered to construct this unified system of multi-layer regulation. The complex relationships between different layers may be approximated by the inter-connected nodes with non-linear activation functions of the neural network. In this paper, we focus on one middle layer of intermediate omics data in the neural network such as gene expression levels.

We have proposed a Bayesian neural network to extend mixed models to multi-layer neural networks to capture the non-linear relationships between genotypes and phenotypes for both genomic prediction and GWAS (Zhao *et al*. 2021). This model, however, is not able to incorporate intermediate omics data. In our proposed neural network named NN-LMM in this paper, omics data are incorporated into the middle layer. An example of the framework of NN-LMM incorporating intermediate omics data is shown in Figure 2, where the nodes in the middle layer represent both observed and unobserved intermediate omics features that are affected by upstream genotypes and regulate the downstream phenotypes. Linear relationships are assumed between genotypes and omics features in the middle layer, such that pedigree-based BLUP (Henderson 1975; Mrode 2014), GBLUP (Habier *et al*. 2007; VanRaden 2008; Hayes *et al*. 2009b), Bayesian Alphabet (Meuwissen *et al*. 2001; Kizilkaya *et al*. 2010; Habier *et al*. 2011; Park and Casella 2008; Gianola and Fernando 2019; Erbe *et al*. 2012; Moser *et al*. 2015), single-step GBLUP (Misztal *et al*. 2009; Aguilar *et al*. 2010), single-step Bayesian Alphabet (Fernando *et al*. 2014) and other mixed models are employed to sample marker effects or genetic values on intermediate omics features. Nonlinear relationships are assumed between intermediate omics features and the phenotype through the activation function in the neural network such as the sigmoid function. Unobserved intermediate omics features will remain to be hidden nodes that will be sampled, thus NN-LMM allows various missing patterns of omics data. For example, in Figure 2, for an individual, the gene expression levels of the first two genes are 0.9 and 0.1, respectively, and the gene expression level of the last gene is missing to be sampled. The missing patterns of gene expression levels can be different for different individuals. Our multi-layer neural network method here can be considered as an extension to conventional mixed models, where the relationship between the first layer of genotypes and the middle layer of omics features can be modeled by mixed models. Here we name our Bayesian neural network specifically “NN-GBLUP”, “NN-BayesA”, “NN-BayesB”, and “NN-BayesC”, when corresponding mixed models (GBLUP, BayesA, BayeB, and BayesC, respectively) are used. In this paper, we will present our model, study its performance, and compare it to the single-step approach in Christensen *et al*. (2021).

**Figure 2.**
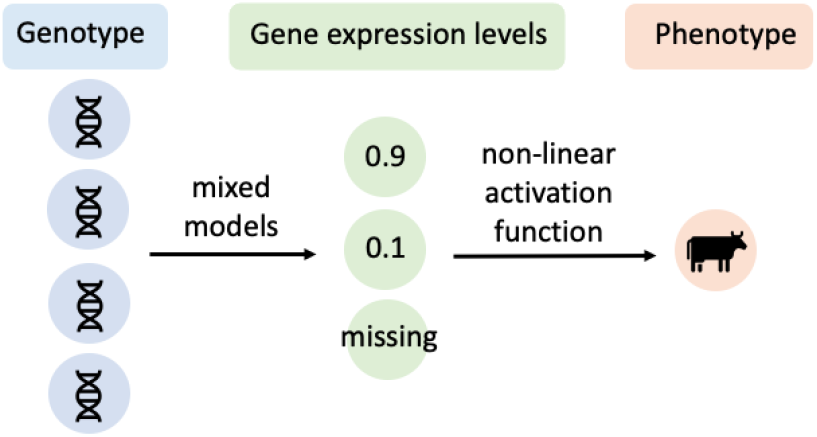
Framework of NN-LMM incorporating intermediate omics data such as gene expression levels. Genotypes affect the gene expression levels, then gene expression levels regulate the phenotypes. Linear mixed models can be applied to sample marker effects or genetic values on gene expression levels, and the nonlinear activation function in neural networks will be used to capture the complex nonlinear relationships between gene expression levels and phenotypes. For an individual, the gene expression levels of the first two genes are 0.9 and 0.1, respectively, and the gene expression of the last gene is missing to be sampled. Individuals can have different missing gene expression levels.

## Materials and methods

A detailed NN-LMM model incorporating intermediate omics features is shown in Figure 3. For *i*th individual, each node in the input layer represents a single-nucleotide polymorphism (SNP) and there are in total *l*_0_ SNPs (i.e., *x*_*i*,1_,…, *x_i_,l*_0_). There are *l*_1_ nodes in the middle layer representing *l*_1_ intermediate omics features (i.e., *z*_*i*,1_,…, *z_i_,l*_1_). Some omics features maybe missing (e.g., orange colored nodes), and different individuals can have different missing omics features. We will use *z_no_* to denote a missing omics feature. The relationship between SNPs and an intermediate omics feature is linear, and priors in conventional mixed models will be used to sample marker effects (i.e., weights between input and middle layers) or genetic values on intermediate omics features. The relationship between intermediate omics features and the phenotype is non-linear, which is approximated by the non-linear activation function of the neural network *g*(·), e.g., the sigmoid function. In NN-LMM, Markov chain Monte Carlo (MCMC) approaches are used to infer unknowns. Below we represent NN-LMM as hierarchical Bayesian regression models.

**Figure 3.**
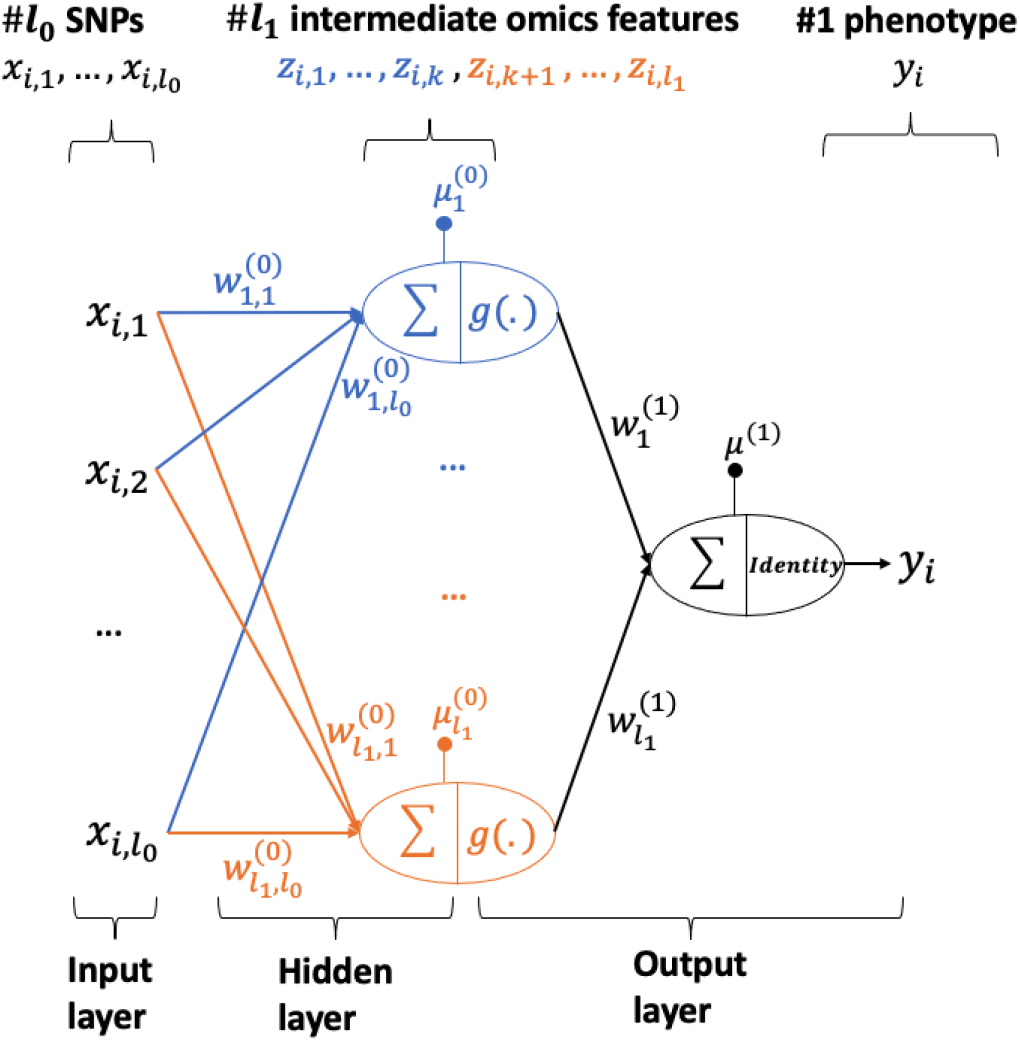
A detailed framework of NN-LMM incorporating intermediate omics data. For *i*th individual, the relationship between SNPs (*x_i,m_*, where *m* = 1,…, *l*_0_) and intermediate omics features (*z_i,j_*, where *j* = 1,…, *l*_1_) is linear, such that linear mixed models are applied to sample marker effects 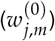 or genetic values of omics features. Non-linear activation function *g*(·) in the neural networks is used to capture the non-linear relationship between intermediate omics features and phenotypes.

### From middle layer (intermediate omics features) to output layer (phenotypes): non-linear activation function

Given all intermediate omics features (observed or sampled), the phenotype of individual *i* is modeled as

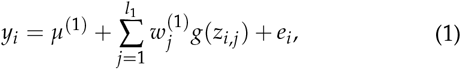

where *y_i_* is the phenotype for individual *i*, *μ*^(1)^ is the overall mean, *z_i,j_* is the *j*th omics feature for individual *i*, *g*(·) is the activation function in neural networks, 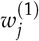 is the effect of *g*(*z_i,j_*) on *y_i_*, and *e_i_* is the random residual. The overall mean *μ*^(1)^ is assigned to a flat prior. The prior of neural network weights 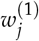 is a normal distribution with null mean and unknown variance 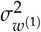, i.e., 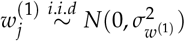. A scaled inverse chi-squared distribution is assigned as the prior for 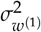, i.e., 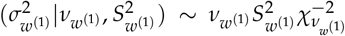. The prior for *e_i_* is a normal distribution with null mean and unknown variance 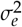, i.e., 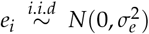. A scaled inverse chi-squared distribution is assigned as the prior for 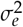, i.e., 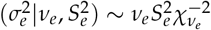.

### From input layer (genotypes) to middle layer (intermediate omics features): mixed models

Given all intermediate omics features (observed or sampled), for *i*th individual, the relationship between the *j*th intermediate omics feature and genotypes can be written as a single-trait mixed model (e.g., Bayesian Alphabet) as:

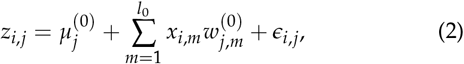

where *z_i,j_* is the *j*th (with *j* = 1,…, *l*_1_) intermediate omics feature for individual *i*, 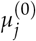 is the overall mean for *j*th intermediate omics feature, *x_i,m_* is the genotype covariate at locus *m* (with *m* = 1,…, *l*_0_) for individual *i* (coded as 0,1,2), 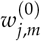 is the marker effects of locus *m* on *j*th intermediate omics feature (i.e., the weight between *m*th node of the input layer and *j*th node of the middle layer), and *ϵ_i,j_* is the random residual of *i*th individual on *j*th intermediate omics feature. Besides Bayesian Alphabet (Meuwissen *et al*. 2001; Kizilkaya *et al*. 2010; Habier *et al*. 2011; Park and Casella 2008; Gianola and Fernando 2019; Erbe *et al*. 2012; Moser *et al*. 2015), the pedigree-based BLUP (Henderson 1975; Mrode 2014), GBLUP (Habier *et al*. 2007; VanRaden 2008; Hayes *et al*. 2009b), single-step GBLUP (Misztal *et al*. 2009; Aguilar *et al*. 2010), or single-step Bayesian Alphabet (Fernando *et al*. 2014) models can also be used to model the relationship between the input and middle layers. The overall mean 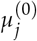 is assigned to a flat prior. Conditional on 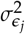, the residuals, *ϵ_ij_*, have independently and identically distributed normal priors with null means and variance 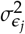, which itself is assumed to have an scaled inverse chi-squared distribution.

Multi-threaded parallelism (Bezanson *et al*. 2017) was implemented to employ multiple single-trait mixed models in parallel at each iteration. When a relatively small number of omics features is used, it is computational feasible to use a multi-trait mixed model to sample marker effects on omics features, e.g., multi-trait BayesC (Cheng *et al*. 2018b). Multi-trait models for pedigree-based BLUP (Henderson and Quaas 1976), GBLUP (Calus and Veerkamp 2011), and single-step methods can also be applied to model the relationship between the first and middle layers.

### Sample missing omics data by Hamiltonian Monte Carlo

For each missing omics feature of *i*th individual (e.g., *z_i,no_*), it will be treated as an unobserved intermediate trait to be sampled by Hamiltonian Monte Carlo (HMC) (Betancourt 2018). HMC will sample the missing omics feature *z_i,no_* from its full conditional distributions.

In HMC, each unknown parameter is paired with a “momentum” variable *ϕ_i,no_*. The HMC constructs the Markov chain by a series of iterations. Following notations in Gelman *et al*. (2013), there are three steps in each iteration of the HMC:

1. updating the momentum variable independently of the current values of the paired parameter, i.e., *ϕ_i,no_* ~ *N*(0, *m*).
2. updating (*z_i,no_, ϕ_i,no_* via *L* leapfrog steps. In each leapfrog step, *z_i,no_* and *ϕ_i,no_* are updated dependently and scaled by *t*. The leapfrog step below is repeated *L* times:

a. 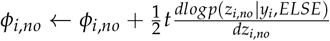;
b. *z_i,no_* ← *z_i,no_* + *tm*^-1^ *ϕ_i,no_*;
c. 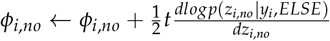. The resulting state at the end of *L* repetitions will be denoted as 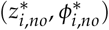.
3. calculating the acceptance rate, *r,* such that the resulting state will be accepted with probability *min*(1, *r*).

As shown above, the gradient of the log full conditional posterior distribution of *z_i_*_,*no*_ is required in HMC, which is:

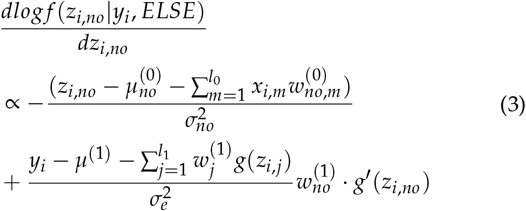

In our analyses, 10 leapfrog steps were used in each HMC iteration, i.e., *L* = 10, *m* was 1, and the scale parameter *t* was 0.1. A more detailed derivation and the full conditional distributions of other parameters of interest in the Gibbs sampler are given in the Appendix.

The estimated breeding value is calculated as 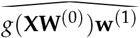, where 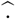 denotes the point estimate of parameters of interest. In NN-LMM, the posterior means are used as the point estimates of parameters of interest.

### Compared to an approach of two mixed model equation system in Christensen et al. (2021)

Christensen *et al*. (2021) proposed a method to include intermediate omics features for genetic evaluation, in which a system of two mixed model equations is required. Using notations in this paper, mixed models in Christensen *et al*. (2021) can be written as:

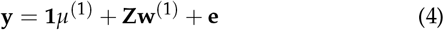

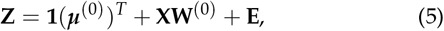

which are equivalent to equation (1) and equation (2) when *g*(·) is a linear activation function.

In equation (4) describing the relationship between phenotypes and intermediate omics features, **y** is the vector of phenotypes, where *y_i_* is the phenotype for individual *i, μ*^(1)^ is the overall mean of phenotypes, **Z** is the matrix of intermediate omics features, where *z_ij_* is the *j*th intermediate omics feature for individual *i*, **w**^(1)^ is the vector of effects of intermediate omics features on phenotypes, where 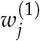 is the effect of *j*th omics feature, **e** is the vector of residuals, where *e_i_* is the residual for *i*th individual. An additional polygenetic effect whose covariance matrix is defined by the pedigree or/and genotypes is also included in equation (4), and this is ignored here for simplicity.

In equation (5) describing the relationship between intermediate omics features and genotypes, ***μ***^(0)^ is the vector of overall means of intermediate omics features, where 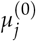 is the overall mean for *j*th intermediate omics feature, **X** is the genotype covariate matrix, where *x_i,m_* is the genotype covariate at locus *m* of *i*th individual, 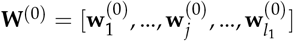 is the matrix of marker effects on all intermediate omics features, where 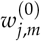 is the marker effects of locus *m* on *j*th intermediate omics feature. **E** is a matrix of residual of intermediate omics features, where *E_i,j_* =*ϵ_i,j_* is the random residual of *i*th individual on *j*th intermediate omics feature.

The genomic breeding values on phenotypes can be calculated as the sum of weighted breeding value from each omics feature, i.e., (**XW**^(0)^)**w**^(1)^, where **XW**^(0)^ is regarded as the breeding values of omics features. In Christensen *et al*. (2021), a matrix of breeding values **G** = [**g**_1_,…, **g***_j_*,…, **g***l*_1_] of omics features, instead of 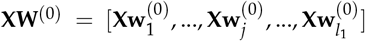, is used, which is similar to expressing SNP-BLUP as GBLUP, and these two models are equivalent in terms of breeding value prediction. Further, breeding values **g**_*j*_ in equation (5) are fitted as random effects, whose covariance matrix is defined by the relationship matrix **H** computed from genotypes or/and pedigree, i.e., 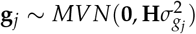.

When all omics features are measured on all individuals, the system of two mixed model equations in Christensen *et al*. (2021) can be regarded as a special case of NN-LMM with a linear activation function between the middle layer and the output layer and a normal prior for marker effects on omics features. However, by extending the mixed model to multi-layer neural networks with non-linear activation functions, NN-LMM may capture non-linear relationships between intermediate omics features and phenotypes. Moreover, NN-LMM allows various priors for marker effects.

#### single-step approach for incomplete omics data

When some individuals do not have an observation on any omics feature, Christensen *et al*. (2021) proposed an approach that is similar to the conventional single-step method. In conventional single-step method (Legarra *et al*. 2009; Christensen and Lund 2010; Legarra *et al*. 2014), when the genotype data are completely missing for some individuals in the pedigree, the phenotypes of these individuals are incorporated by modelling the covariances of their breeding values with those of the genotyped individuals through pedigree relationships. In Christensen *et al*. (2021), a similar single-step approach is used to incorporate the phenotypes of individuals with missing omics data by modelling the covariances of their omics values with those of the omics-typed individuals through genomic relationships, where the omics value of an individual is computed as the sum of omics contributions on phenotypes in equation (4), i.e., **u** = **Zw**^(1)^. In detail, when all omics data are observed on all individuals, **u** can be considered as random effects whose covariance matrix is 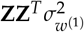. When all omics features are missing for some individuals, however, **Z** for all individuals are not observed. Christensen *et al*. (2021) proposed that the covariance matrix for the sum of omics contributions on phenotypes for all individuals, i.e., **u** = **Zw**^(1)^, can be computed by combining information in 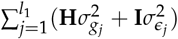 for all individuals and that in the omics relationship matrix 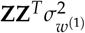 for individuals with omics data.

A potential issue with above single-step approach is that the residual part 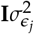 is included along with the genomic and/or pedigree relationship matrix 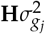. Thus, when many individuals have no omics measures, this residual part may be dominant. As we will show below, compared to this single-step approach, NN-LMM provides a more straightforward approach to analyze datasets when some individuals have no omics measures, and gives equivalent or higher prediction accuracy even when the underlying relationships between intermediate omics features and phenotypes are linear. Also, NN-LMM can handle various missing patterns in omics data, whereas the single-step approach in Christensen *et al*. (2021) only works when all omics features are not measured on some individuals.

### Data Analysis

#### Linear system

To compare the prediction performance of NN-LMM to the single-step approach in Christensen *et al*. (2021), a linear activation function was used in NN-LMM, consistent with the assumption in the single-step approach that the relationship between intermediate omics features and phenotypes is linear.

The goal of genetic evaluation is to accurately predict breeding values, rather than phenotypes, and it is not straightforward to validate the prediction of breeding values using real datasets. Thus, simulated data from Christensen *et al*. (2021) were used. Note that in Christensen *et al*. (2021), an polygenic effect whose covariance matrix is defined by the pedigree or/and genotypes is also included in equation (4). This part is ignored here for simplicity, and was subtracted from the simulated phenotypes. Two patterns of missing omics data were considered: missing omics pattern (i) as in Christensen *et al*. (2021): all omics features are not measured on some individuals; missing omics pattern (ii): for each omics feature, some random individuals have no omics data. Missing omics pattern (i) is a special case of missing omics pattern (ii). The single-step approach only works with the scenario (i), while NN-LMM allows both scenarios.

The simulated data in Christensen *et al*. (2021) contained 21,100 individuals from 11 generations. We randomly sampled 5% individuals from each generation to have a subset of 1,055 individuals (i.e., 55 individuals in the first generation, and 100 individuals in each of the later generations). Following Christensen *et al*. (2021), individuals in the last generation were used as the testing dataset (i.e., 100 individuals), whose omics features were observed but phenotypes were unknown, and the remaining individuals (i.e., 955 individuals) were used for training. The genotypic data consists of 15,000 SNP markers observed for all individuals, and the intermediate omics data consists of 1,200 omics features. Each omics feature was affected by 500 QTLs randomly selected from a set of 5,000 QTLs which were not included in the 15,000 SNP markers, and phenotypes were affected by all 1,200 intermediate omics features. The heritability of each omics feature was 0.61, and the heritability of the phenotypic trait was 0.337. More details about the simulation process are in Christensen *et al*. (2021).

When all individuals have all omics features measured, the performance of NN-LMM and the system of two mixed model equations in Christensen *et al*. (2021) were compared using 20 replicates. Different proportions of missing omics data in the training dataset were considered, including 0%, 10%, 30%, 50%, 70%, 80%, 90%, 95%, 99%, where 0% denotes a scenario where all omics features are measured on all individuals. For each scenario, 20 replicates were used. The GBLUP model in conventional genomic evaluation, where no omics data are available, was used as the baseline for comparison.

The prediction accuracy was calculated as the Pearson correlation between the true breeding values **XW**^(0)^**w**^(1)^ and the estimated breeding values (i.e., 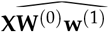 in NN-LMM and 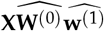 in the single-step approach) for individuals in the testing datasets, where 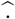 denotes the point estimate of parameters of interest. In NN-LMM with linear activation function, a number of 5,000 MCMC iterations was applied to ensure the convergence.

#### Nonlinear system

Studies have shown that the relationship between intermediate omics features and phenotypes may be non-linear (Kitano 2002; Green *et al*. 2017; Devijver *et al*. 2017; Green *et al*. 2019). One example is that Green *et al*. (2017) used a von Bertalanffy growth curve, which is a generalised logistic function, to approximate the relationship between gene expression levels and a quantitative trait. Using data in Christensen *et al*. (2021), we simulated nonlinear relationships between intermediate omics features and phenotypes. In detail, the sigmoid non-linear transformation was applied to the omics data, and the phenotypes were affected by the nonlinear-transformed omics data as in equation (1). Same heritability, variance components were used as in the above linear system, as well as the number of omics features (i.e., 1200) and the number of randomly selected QTLs affecting each omics feature (i.e., 500). The QTLs were also not included in the SNP markers.

The performance of NN-LMM were studied using different activation functions in neural networks, including a linear function (i.e., the identity function) and a non-linear function (i.e., the sigmoid function). Both missing omics patterns (i) and (ii) were considered, and different proportions of missing omics data were tested. 10 replicates were applied in each scenario. The prediction accuracy was calculated as the Pearson correlation between the true breeding values *g*(**XW**^(0)^)**w**^(1)^ and the estimated breeding values 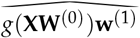 for individuals in the testing datasets. MCMC chains of length 5,000 and 20,000 were applied to NN-GBLUP with sigmoid and linear activation functions, respectively, to ensure the convergence.

## Results

### Linear system

To compare the prediction performance of NN-LMM to the single-step approach in Christensen *et al*. (2021), a linear activation function was used in NN-LMM, consistent with the assumption in the single-step approach that the relationship between intermediate omics features and phenotypes is linear.

When all omics features are measured on all individuals, the system of two mixed model equations in Christensen *et al*. (2021) can be regarded as a special case of NN-LMM with a linear activation function between the middle layer and the output layer and a normal prior for marker effects on omics features. Following Christensen *et al*. (2021), variance components were treated as known for both methods to be the values used in the simulation. NN-GBLUP had similar prediction accuracies as the system of two mixed model equations in Christensen *et al*. (2021) for all 20 replicates (correlation *r* = 0.999).

Results for missing omics patterns (i) and (ii) were shown in Figure 4. Overall, the prediction accuracy decreased when the proportion of missing omics data increased. For missing omics pattern (i), when a small proportion of individuals had no omics data, NN-GBLUP (red solid line) had similar prediction performance as the single-step approach in Christensen *et al*. (2021) (blue solid line). However, when a large proportion of individuals had no omics data (e.g., >80%), NN-GBLUP had significantly higher prediction accuracies (pairwise t-test *P*-value < 0.005). When >90% individuals had no omics data, the single-step approach performed even worse than the baseline (black dashed line), which was a conventional GBLUP model where no omics data were used. For missing omics pattern (ii), the prediction accuracy of NN-GBLUP (red dashed line) decreased with larger proportion of missing omics data, and eventually close to the baseline, whereas the single-step approach did not work for this scenario.

**Figure 4.**
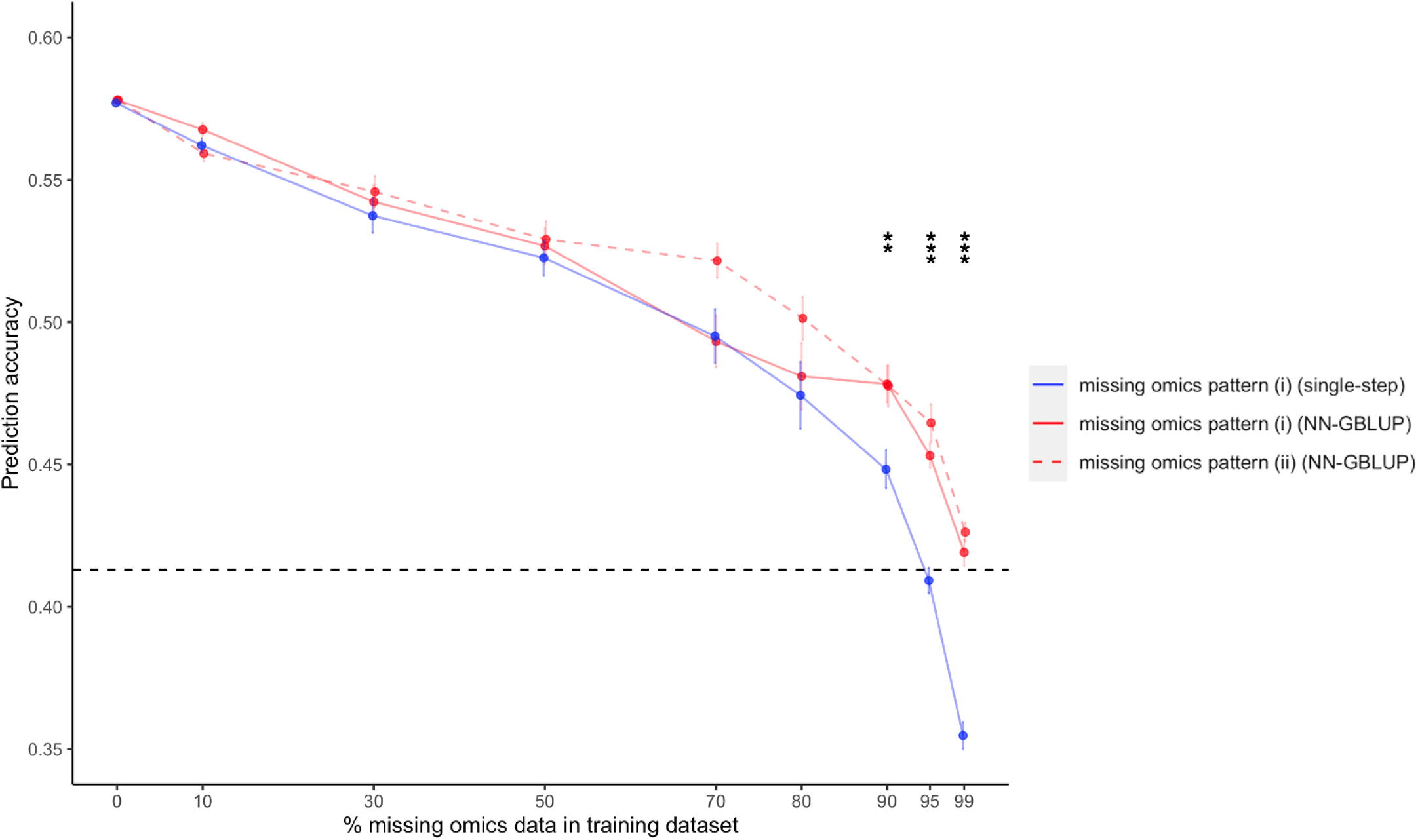
Prediction accuracies of NN-GBLUP with the linear activation function and the single-step approach in Christensen *et al*. (2021). Different proportions of missing omics data in the training dataset were considered, including 0%, 10%, 30%, 50%, 70%, 80%, 90%, 95%, 99%. There were two missing omics patterns: missing omics pattern (i): all omics features were not measured on some individuals, and missing omics pattern (ii): for each omics feature, some random individuals had no omics measures. Missing omics pattern (i) is a special case of pattern (ii), and the single-step approach only works with the pattern (i). The horizontal black dashed linear represents the conventional GBLUP model when no omics data were available, and it was used as the baseline for both methods. Each dot represents the mean of prediction accuracies from 20 replications, and the vertical bar is the mean ± its standard error. The asterisk symbol indicated that for missing omics pattern (i), NN-GBLUP had significantly higher prediction accuracy than the single-step approach under the t-test with a significance level of 0.005 (**) or lower (* * *).

### Nonlinear system

Compared to the system of two linear models (Weishaar *et al*. 2020; Christensen *et al*. 2021; Gamazon *et al*. 2015; Gusev *et al*. 2016; Wainberg *et al*. 2019), NN-LMM allows nonlinear relationships between intermediate omics features and phenotypes. Under the nonlinear system, the underlying relationship between intermediate omics features and phenotypes was nonlinear for the simulated datasets. The prediction performance of NN-GBLUP with a linear function was compared to NN-GBLUP with a nonlinear sigmoid activation function. All variance components were sampled.

For different proportions of missing omics data under both missing omics patterns, using the nonlinear activation function in NN-LMM was significantly better than using the linear activation function. The results for the proportion of 50% missing omics data were shown in Figure 5 (pairwise t-test *P*-value < 0.0001).

**Figure 5.**
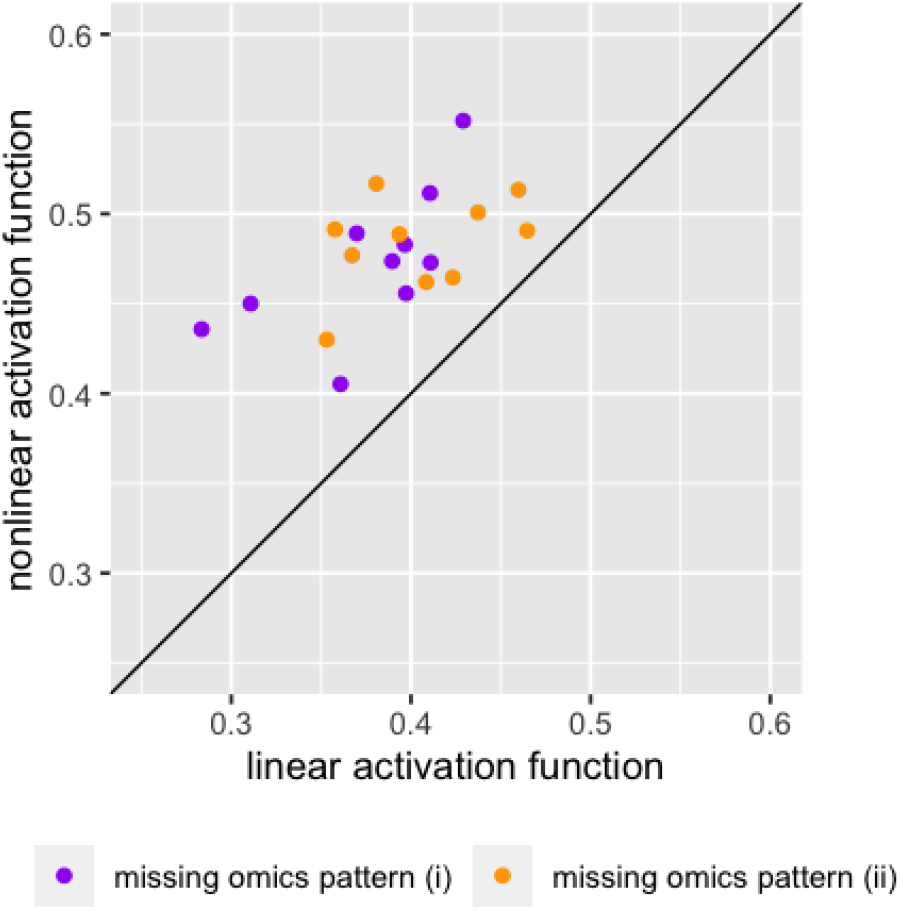
The prediction performance of NN-GBLUP with the linear activation function versus NN-GBLUP with the nonlinear sigmoid activation function, when there were 50% missing omics data in the training dataset. Missing omics patterns (i) and (ii) were distinguished by color, and 10 replicates were applied for each pattern. The diagonal line was used for reference such that a dot above the line represents a replicate with higher prediction accuracy for the nonlinear sigmoid activation function.

### Effects of different priors in NN-LMM

NN-LMM allows various priors from conventional mixed models to model the relationship between the genotypes and intermediate omics features. Here the performance of NN-LMM with a GBLUP prior (i.e, NN-GBLUP) and NN-LMM with a BayesC prior (i.e., NN-BayesC) were compared, and linear activation functions were applied. All variance components were sampled.

NN-GBLUP had similar prediction accuracies as NN-BayesC for the above simulated datasets. This may be due to the relatively small sample size (*n* = 1, 055) compared to the number of SNPs (*p* = 15, 000). Thus, we selected 1,000 SNPs evenly from all 15,000 SNPs, and 250 SNPs were randomly selected as QTLs from this 1,000 SNP panel. 20 intermediate omics features were simulated, where each omics feature was affected by 50 QTLs randomly selected from the 250 QTLs. QTLs were included in the SNP panel. Heritability and variance components were the same as before.

For different proportions of missing omics data, NN-BayesC had significantly higher prediction accuracy than NN-GBLUP under both missing omics patterns (pairwise t-test *P*-value < 0.0001). The results for the proportion of 50% missing omics data were shown in Figure 6.

**Figure 6.**
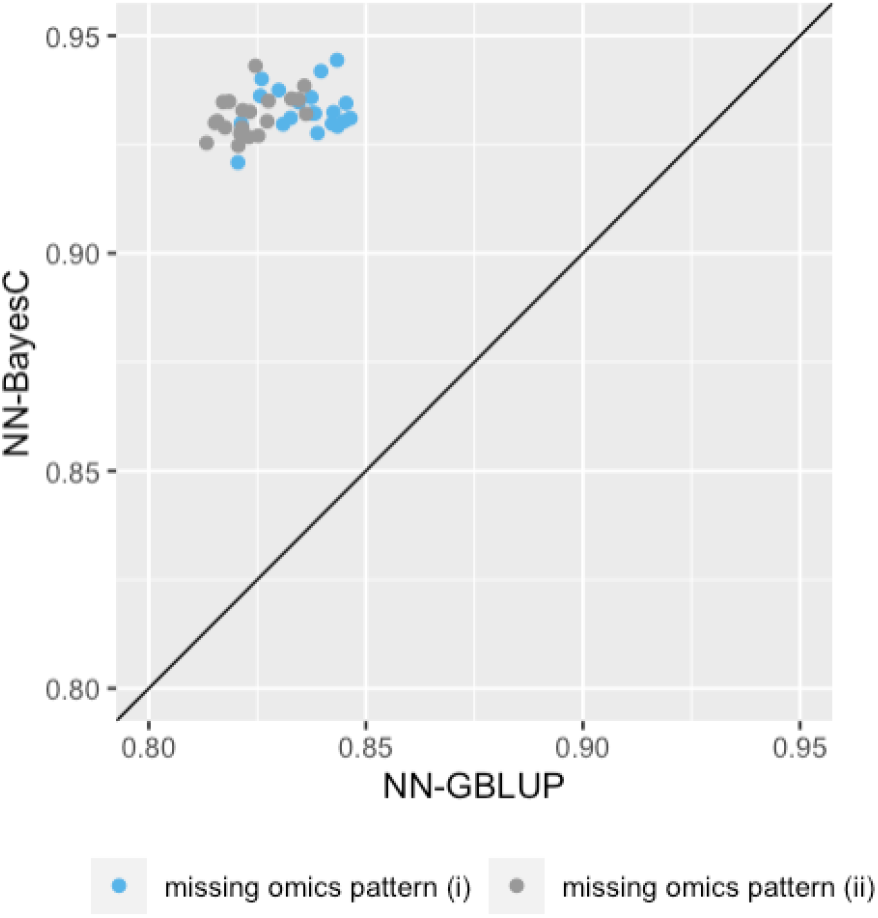
The prediction performance of NN-GBLUP versus NN-BayesC, when there were 50% missing omics data in the training dataset. Linear activation function was used. Missing omics patterns (i) and (ii) were distinguished by color, and 20 replicates were applied for each pattern. The diagonal line was used for reference such that a dot above the line represents a replicate with higher prediction accuracy for the NN-BayesC.

## Discussion

Using omics data only, especially the gene expression levels, to predict the phenotype is not new (e.g., Golub *et al*. (1999); Riedelsheimer *et al*. (2012); Li *et al*. (2019); Guo *et al*. (2016)). However, such methods for the prediction of phenotypic values do not lead in themselves for genetic improvement (i.e., better estimation of breeding values). In association studies, to incorporate omics data as intermediate traits between genotypes and phenotypes (Ritchie *et al*. 2015; Gamazon *et al*. 2015; Gusev *et al*. 2016; Wainberg *et al*. 2019), two linear models were used, where one model describes how genotypes affect omics features, and another describes how omics features affect phenotypes. Recently, such a system of two linear models has also been developed for genomic evaluation (Weishaar *et al*. 2020; Christensen *et al*. 2021), and further extended for genomic prediction using incomplete omics data when some individuals had no omics measures (Christensen *et al*. 2021).

In this paper, we proposed a new method named NN-LMM to extend linear mixed models to multi-layer neural networks for genomic prediction with intermediate omics features. NN-LMM models the unified system of multi-layer regulations from genotypes to intermediate omics features, then to the phenotype, such that the upstream genotypes affect the intermediate omics features, then omics features regulate the downstream phenotypes. Compared to other methods, NN-LMM provides a more flexible and robust framework to incorporate intermediate omics features. First, NN-LMM allows various patterns of missing omics data, for example, individuals can have different missing omics features. Second, NN-LMM allows nonlinear relationships between intermediate omics features and phenotypes, and the non-linear relationships are approximated by activation functions in neural networks. Third, various linear mixed models can be used to model the relationship between the genotypes and omics features. NN-LMM has been implemented in an open-source package called “JWAS” (Cheng *et al*. 2018a).

In simulation analysis, NN-LMM had significantly higher prediction accuracy than the single-step approach in Christensen *et al*. (2021) when a large proportion of individuals had no omics data. As shown in Table 1, however, incorporating those individuals with no omics data, either using the single-step approach or NN-LMM, was better than simply deleting them from the dataset.

**Table 1.** Comparison of the prediction performance between the strategy of deleting individuals with no omics data and incorporating those individuals via the single-step approach or NN-LMM.

| Method | % missing omics data in training dataset |  |  |  |  |  |  |  |  |
| --- | --- | --- | --- | --- | --- | --- | --- | --- | --- |
|  | 0% | 10% | 30% | 50% | 70% | 80% | 90% | 95% | 99% |
| delete individuals with no omics data | 0.577<sup>a</sup> | 0.564 | 0.526 | 0.487 | 0.419 | 0.357 | 0.291 | 0.218 | 0.096 |
| single-step approach | 0.577 | 0.562 | 0.537 | 0.523 | 0.495 | 0.474 | 0.448 | 0.409 | 0.355 |
| NN-GBLUP | 0.578 | 0.568 | 0.542 | 0.527 | 0.493 | 0.481 | 0.478 | 0.453 | 0.419 |

<sup>a</sup> The prediction accuracy was presented as the mean of prediction accuracies from 20 replications.

NN-LMM allows nonlinear relationships between intermediate omics features and phenotypes. In our simulation analysis, when the underlying relationship between intermediate omics features and phenotypes was nonlinear, using the nonlinear activation function in NN-LMM had significantly better performance than using the linear activation function. Given the observations that the relationships between intermediate omics features and the phenotypes might be nonlinear (Kitano 2002; Green *et al*. 2017; Devijver *et al*. 2017; Green *et al*. 2019), NN-LMM may be a more biological realistic approach than other system of linear models.

However, one issue with the current implementation of NN-LMM is computation. To sample marker effects on omics features, a naive multi-threaded parallelism (Bezanson *et al*. 2017) has been implemented to employ multiple single-trait mixed models in parallel at each MCMC iteration. Thus, ideally, with thousands of computer processors, the running time to sample marker effects on thousands of omics features equals that of one omics feature (i.e., one single-trait mixed model). However, due to the hardware limitation (e.g., on a personal laptop), this parallelisation was usually only a few times faster than without parallel computing. In practice, it took about 10 hours on a personal laptop to run 5,000 iterations for a dataset with 1,055 individuals, 15,000 SNPs and 1,200 omics features. Whereas solving the system of two mixed model equations without estimating variance components in Christensen *et al*. (2021) only required a few minutes for such a dataset. To improve the computation performance of NN-LMM in the future, parallel computing strategies, e.g., the strategy in Zhao *et al*. (2020), needs to be further studied.

In NN-LMM, unknowns were sampled from their full conditional posterior distributions, and the posterior means were used as the point estimates of parameters of interest. Thus, the estimated breeding value is calculated as 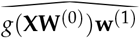, where 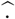 denotes the point estimate of parameters of interest. When the relationship between omics features and phenotypes is linear, the estimated breeding value in NN-LMM is 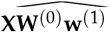, whereas the estimated breeding value in Christensen *et al*. (2021) is 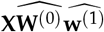, assuming **W**^(0)^ and **w**^(1)^ are independent. This assumption of independence may affect the model performance. Note that the goal of genetic evaluation is to accurately predict breeding values, rather than phenotypes. Thus, caution is needed when comparing NN-LMM to methods for phenotypic prediction.

Another approach for genomic prediction using omics features is to include both target phenotype and omics features as correlated traits in a multi-trait genetic model (Hayes *et al*. 2017). However, it would be computational infeasible to include high-dimensional omics features, such as the expression levels of thousand of genes, in a multi-trait model. Runcie *et al*. (2021) recently proposed a linear mixed model for genomic predictions with thousands of traits. However, it is difficult to model directional regulatory cascades in a multi-trait model framework when considering multiple layers of omics data, which is straightforward for NN-LMM.

## Data Availability Statement

Simulated datasets used in the analysis are publicly available in Christensen *et al*. (2021). All scripts are available at https://github.com/zhaotianjing/NN-LMM. The authors state that all data necessary for confirming the conclusions presented in the article are represented fully within the article.

## Acknowledgements

This work was supported by the United States Department of Agriculture, Agriculture and Food Research Initiative National Institute of Food and Agriculture Competitive Grant No. 2018-67015-27957 and No. 2021-67015-33412.

## Appendix MCMC in NN-LMM

### sampling effects of omics features on phenotypes

In NN-LMM, the effects of intermediate omics features on phenotypes are weights between middle layer and output layer, 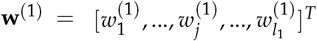, with prior 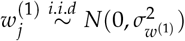. The full conditional posterior distribution of **w**^(1)^ is a multivariate normal distribution with mean

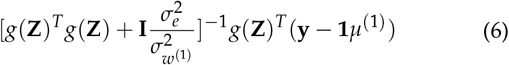

and covariance matrix 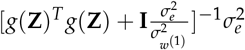.

### sampling the overall mean of phenotypes

The overall mean of phenotypes is *μ*^(1)^ in equation 1 with a flat prior. The full conditional posterior distribution of *μ*^(1)^ is a normal distribution with mean (**1***^T^* **1**)^-1^**1***^T^*(**y** – *g*(**Z**)**w**^(1)^) and variance 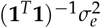.

### sampling missing omics features

The gradient of the log full conditional posterior distribution of the missing omics feature for individual *i*, i.e., *z_i,no_*, is derived below. The full conditional posterior distribution of *z_i,no_* can be expressed as:

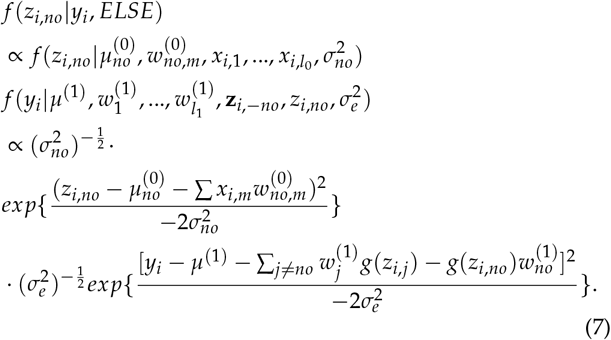

Then, the log full conditional posterior distribution of *z_i,no_* is:

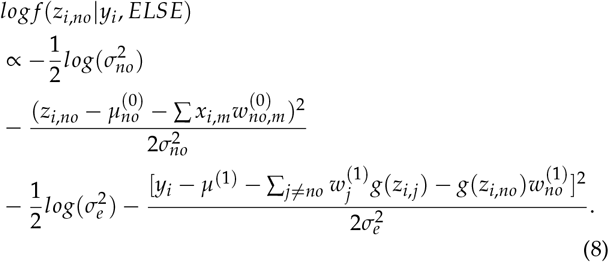

Thus, the gradient of the log-full conditional posterior distribution of *z_i,no_* is:

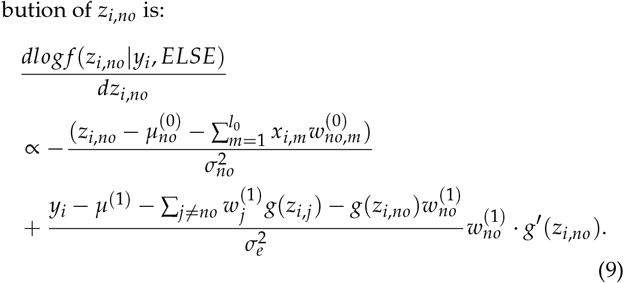

## References

Aguilar, I., I. Misztal, D. Johnson, A. Legarra, S. Tsuruta, et al., 2010 Hot topic: A unified approach to utilize phenotypic, full pedigree, and genomic information for genetic evaluation of holstein final score. Journal of dairy science 93: 743–752.

Atwell, S., Y. S. Huang, B. J. Vilhjálmsson, G. Willems, M. Horton, et al., 2010 Genome-wide association study of 107 phenotypes in arabidopsis thaliana inbred lines. Nature 465: 627–631.

Betancourt, M., 2018 A conceptual introduction to hamiltonian monte carlo.

Bezanson, J., A. Edelman, S. Karpinski, and V. B. Shah, 2017 Julia: A fresh approach to numerical computing. SIAM review 59: 65–98.

Calus, M. P. and R. F. Veerkamp, 2011 Accuracy of multi-trait genomic selection using different methods. Genetics Selection Evolution 43: 1–14.

Cheng, H., R. Fernando, and D. Garrick, 2018a Jwas: Julia implementation of whole-genome analysis software. In Proceedings of the world congress on genetics applied to livestock production, volume 11, p. 859.

Cheng, H., K. Kizilkaya, J. Zeng, D. Garrick, and R. Fernando, 2018b Genomic prediction from multiple-trait bayesian regression methods using mixture priors. Genetics 209: 89–103.

Christensen, O. F., V. Börner, L. Varona, and A. Legarra, 2021 Genetic evaluation including intermediate omics features. Genetics iyab130.

Christensen, O. F. and M. S. Lund, 2010 Genomic prediction when some animals are not genotyped. Genetics Selection Evolution 42: 1–8.

Devijver, E., M. Gallopin, and E. Perthame, 2017 Nonlinear network-based quantitative trait prediction from transcriptomic data. arXiv preprint arXiv:1701.07899.

Erbe, M., B. Hayes, L. Matukumalli, S. Goswami, P. Bowman, et al., 2012 Improving accuracy of genomic predictions within and between dairy cattle breeds with imputed high-density single nucleotide polymorphism panels. Journal of dairy science 95: 4114–4129.

Fernando, R. L., J. C. Dekkers, and D. J. Garrick, 2014 A class of bayesian methods to combine large numbers of genotyped and non-genotyped animals for whole-genome analyses. Genetics Selection Evolution 46: 1–13.

Gamazon, E. R., H. E. Wheeler, K. P. Shah, S. V. Mozaffari, K. Aquino-Michaels, et al., 2015 A gene-based association method for mapping traits using reference transcriptome data. Nature genetics 47: 1091–1098.

Gelman, A., J. B. Carlin, H. S. Stern, D. B. Dunson, A. Vehtari, et al., 2013 Bayesian data analysis. CRC press.

Gianola, D. and R. L. Fernando, 2019 A Multiple-Trait Bayesian Lasso for Genome-Enabled Analysis and Prediction of Complex Traits. Genetics 214: genetics.302934.2019–331.

Golub, T. R., D. K. Slonim, P. Tamayo, C. Huard, M. Gaasenbeek, et al., 1999 Molecular classification of cancer: class discovery and class prediction by gene expression monitoring. science 286: 531–537.

Green, R. M., J. L. Fish, N. M. Young, F. J. Smith, B. Roberts, et al., 2017 Developmental nonlinearity drives phenotypic robustness. Nature communications 8: 1–12.

Green, R. M., C. L. Leach, V. M. Diewert, J. D. Aponte, E. J. Schmidt, et al., 2019 Nonlinear gene expression-phenotype relationships contribute to variation and clefting in the a/wysn mouse. Developmental Dynamics 248: 1232–1242.

Guo, Z., M. M. Magwire, C. J. Basten, Z. Xu, and D. Wang, 2016 Evaluation of the utility of gene expression and metabolic information for genomic prediction in maize. Theoretical and applied genetics 129: 2413–2427.

Gusev, A., A. Ko, H. Shi, G. Bhatia, W. Chung, et al., 2016 Integrative approaches for large-scale transcriptome-wide association studies. Nature genetics 48: 245–252.

Habier, D., R. L. Fernando, and J. C. Dekkers, 2007 The impact of genetic relationship information on genome-assisted breeding values. Genetics 177: 2389–2397.

Habier, D., R. L. Fernando, K. Kizilkaya, and D. J. Garrick, 2011 Extension of the bayesian alphabet for genomic selection. BMC bioinformatics 12: 186.

Hayes, B., J. Panozzo, C. Walker, A. Choy, S. Kant, et al., 2017 Accelerating wheat breeding for end-use quality with multi-trait genomic predictions incorporating near infrared and nuclear magnetic resonance-derived phenotypes. Theoretical and applied genetics 130: 2505–2519.

Hayes, B. J., P. J. Bowman, A. J. Chamberlain, and M. E. Goddard, 2009a Invited review: Genomic selection in dairy cattle: Progress and challenges. Journal of dairy science 92: 433–443.

Hayes, B. J., P. M. Visscher, and M. E. Goddard, 2009b Increased accuracy of artificial selection by using the realized relationship matrix. Genetics research 91: 47–60.

Heffner, E. L., M. E. Sorrells, and J.-L. Jannink, 2009 Genomic selection for crop improvement. Crop Science 49: 1–12.

Henderson, C. and R. Quaas, 1976 Multiple trait evaluation using relatives’ records. Journal of animal science 43: 1188–1197.

Henderson, C. R., 1975 Best linear unbiased estimation and prediction under a selection model. Biometrics pp. 423–447.

Hickey, J. M., T. Chiurugwi, I. Mackay, W. Powell, A. Eggen, et al., 2017 Genomic prediction unifies animal and plant breeding programs to form platforms for biological discovery. Nature genetics 49: 1297.

Kitano, H., 2002 Computational systems biology. Nature 420: 206–210.

Kizilkaya, K., R. L. Fernando, and D. J. Garrick, 2010 Genomic prediction of simulated multibreed and purebred performance using observed fifty thousand single nucleotide polymorphism genotypes1. Journal of Animal Science 88: 544–551.

Korte, A. and A. Farlow, 2013 The advantages and limitations of trait analysis with gwas: a review. Plant methods 9: 1–9.

Legarra, A., I. Aguilar, and I. Misztal, 2009 A relationship matrix including full pedigree and genomic information. Journal of dairy science 92: 4656–4663.

Legarra, A., O. F. Christensen, I. Aguilar, and I. Misztal, 2014 Single step, a general approach for genomic selection. Livestock Science 166: 54–65.

Li, Z., N. Gao, J. W. Martini, and H. Simianer, 2019 Integrating gene expression data into genomic prediction. Frontiers in genetics 10: 126.

Meuwissen, T. H. E., B. J. Hayes, and M. E. Goddard, 2001 Predic-tion of total genetic value using genome-wide dense marker maps. Genetics 157: 1819–1829.

Misztal, I., A. Legarra, and o. Aguilar, 2009 Computing procedures for genetic evaluation including phenotypic, full pedigree, and genomic information. Journal of dairy science 92: 4648–4655.

Moser, G., S. H. Lee, B. J. Hayes, M. E. Goddard, N. R. Wray, et al., 2015 Simultaneous discovery, estimation and prediction analysis of complex traits using a bayesian mixture model. PLoS Genet 11: e1004969.

Mrode, R. A., 2014 Linear models for the prediction of animal breeding values. Cabi.

Ozaki, K., Y. Ohnishi, A. Iida, A. Sekine, R. Yamada, et al., 2002 Functional snps in the lymphotoxin-*α* gene that are associated with susceptibility to myocardial infarction. Nature genetics 32: 650–654.

Park, T. and G. Casella, 2008 The bayesian lasso. Journal of the American Statistical Association 103: 681–686.

Qian, J., E. Ray, R. L. Brecha, M. P. Reilly, and A. S. Foulkes, 2019 A likelihood-based approach to transcriptome association analysis. Statistics in medicine 38: 1357–1373.

Riedelsheimer, C., A. Czedik-Eysenberg, C. Grieder, J. Lisec, F Technow, et al., 2012 Genomic and metabolic prediction of complex heterotic traits in hybrid maize. Nature genetics 44: 217–220.

Ritchie, M. D., E. R. Holzinger, R. Li, S. A. Pendergrass, and D. Kim, 2015 Methods of integrating data to uncover genotype–phenotype interactions. Nature Reviews Genetics 16: 85–97.

Runcie, D. E., J. Qu, H. Cheng, and L. Crawford, 2021 Megalmm: Mega-scale linear mixed models for genomic predictions with thousands of traits. Genome biology 22: 1–25.

Sun, Y. V. and Y.-J. Hu, 2016 Integrative analysis of multi-omics data for discovery and functional studies of complex human diseases. Advances in genetics 93: 147–190.

VanRaden, P. M., 2008 Efficient methods to compute genomic predictions. Journal of dairy science 91: 4414–4423.

Visscher, P. M., M. A. Brown, M. I. McCarthy, and J. Yang, 2012 Five years of gwas discovery. The American Journal of Human Genetics 90: 7–24.

Visscher, P. M., N. R. Wray, Q. Zhang, P. Sklar, M. I. McCarthy, et al., 2017 10 years of gwas discovery: biology, function, and translation. The American Journal of Human Genetics 101: 5–22.

Wainberg, M., N. Sinnott-Armstrong, N. Mancuso, A. N. Barbeira, D. A. Knowles, et al., 2019 Opportunities and challenges for transcriptome-wide association studies. Nature genetics 51: 592–599.

Weishaar, R., R. Wellmann, A. Camarinha-Silva, M. Rodehutscord, and J. Bennewitz, 2020 Selecting the hologenome to breed for an improved feed efficiency in pigs—a novel selection index. Journal of Animal Breeding and Genetics 137: 14–22.

Wu, Y., J. Zeng, F. Zhang, Z. Zhu, T. Qi, et al., 2018 Integrative analysis of omics summary data reveals putative mechanisms underlying complex traits. Nature communications 9: 1–14.

Zhao, T., R. Fernando, and H. Cheng, 2021 Interpretable artificial neural networks incorporating Bayesian alphabet models for genome-wide prediction and association studies. G3 Genes | Genomes | Genetics 11, jkab228.

Zhao, T., R. Fernando, D. Garrick, and H. Cheng, 2020 Fast parallelized sampling of Bayesian regression models for whole-genome prediction. Genetics Selection Evolution 52: 1–11.,

